# Blended Genome Exome (BGE) as a Cost Efficient Alternative to Deep Whole Genomes or Arrays

**DOI:** 10.1101/2024.04.03.587209

**Authors:** Matthew DeFelice, Jonna L. Grimsby, Daniel Howrigan, Kai Yuan, Sinéad B. Chapman, Christine Stevens, Samuel DeLuca, Megan Townsend, Joseph Buxbaum, Margaret Pericak-Vance, Shengying Qin, Dan J. Stein, Solomon Teferra, Ramnik J. Xavier, Hailiang Huang, Alicia R. Martin, Benjamin M. Neale

**Author notes:** **Corresponding author:** Matthew DeFelice; The Broad Institute of MIT and Harvard, 320 Charles St. Cambridge, MA 02141, U.S.A. authors contributed equally to this work (Matthew DeFelice, Jonna L. Grimsby). **Email addresses of authors:** DeFelice, Grimsby, Howrigan, Yuan, Chapman, Stevens, DeLuca, Townsend, Buxbaum, Pericak-Vance, Qin, Stein, Teferra, Xavier, Huang, Martin, Neale.

## Abstract

Genomic scientists have long been promised cheaper DNA sequencing, but deep whole genomes are still costly, especially when considered for large cohorts in population-level studies. More affordable options include microarrays + imputation, whole exome sequencing (WES), or low-pass whole genome sequencing (WGS) + imputation. WES + array + imputation has recently been shown to yield 99% of association signals detected by WGS. However, a method free from ascertainment biases of arrays or the need for merging different data types that still benefits from deeper exome coverage to enhance novel coding variant detection does not exist. We developed a new, combined, “Blended Genome Exome” (BGE) in which a whole genome library is generated, an aliquot of that genome is amplified by PCR, the exome regions are selected and enriched, and the genome and exome libraries are combined back into a single tube for sequencing (33% exome, 67% genome). This creates a single CRAM with a low-coverage whole genome (2-3x) combined with a higher coverage exome (30-40x). This BGE can be used for imputing common variants throughout the genome as well as for calling rare coding variants. We tested this new method and observed >99% r^2^ concordance between imputed BGE data and existing 30x WGS data for exome and genome variants. BGE can serve as a useful and cost-efficient alternative sequencing product for genomic researchers, requiring ten-fold less sequencing compared to 30x WGS without the need for complicated harmonization of array and sequencing data.

## Main Text

Due to the current pricing of DNA sequencing, whole genome sequencing (WGS) of large cohorts of samples can be cost-prohibitive at the scales needed to robustly discover enome-wide significant loci. Instead, scientists studying associations between genetic variants and disease have used single nucleotide polymorphism (SNP) genotyping arrays to capture common variants^1^ or deep-coverage whole exome sequencing (WES) to capture rare, functional variants in the 1% coding portion of the genome^2^. Genotyping data from SNP arrays is often bolstered by imputation of missing genotypes by making use of linkage-disequilibrium and a smaller reference panel of sequenced whole genomes. Then, genome-wide association studies use regression analysis to identify genomic regions associated with a trait or disease of interest^3^. A recent analysis of genetic association signal yields from the UK Biobank found that WES + array + imputation can detect 99% of signals detected by WGS, suggesting that it should be favored over costly WGS for association discovery in large sample sets.^4^ However, SNP arrays have been developed primarily based on data from European populations, and this impacts population genetic analyses and utility for studies of more diverse global populations^5^. Merging SNP array data with DNA exome sequencing data together, to capture both rare and common variants, presents further challenges for analysis. Low-depth WGS with imputation has recently been shown to be a cost-effective option for capturing both novel and common variants more accurately than commonly used arrays^6^, but the need for a method that benefits from deeper exome coverage for rare coding variant detection without requiring multiple data sets remains unmet.

To overcome these challenges, and to achieve comparable genetic association yields to 30x WGS with ten-fold less sequencing, we developed a new, combined, “Blended Genome Exome” (BGE) that can serve as an alternative and cost-efficient sequencing product for genomic researchers. In this BGE process (Fig. 1), we construct PCR-free whole genomes, take an aliquot for PCR amplification, select the exome region from the amplified genome through hybridization and capture, blend the exome libraries (33%) back with the PCR-free whole genome libraries (67%) (same sample identification index at the ligation event) into a single tube, and sequence them together. This creates a single CRAM (condensed BAM) file with a low-coverage whole genome (2-3x) combined with a higher coverage exome (30-40x). This can be used both for imputing common variants throughout the genome and calling rare variants in coding regions, at a fraction of the cost of deep-coverage whole genomes with no need for merging separate data sets. Here, we present the first high-throughput, in-process, genome and exome blending method to date that combines 384 exomes and genomes pre-sequencing and results in a single blended CRAM file for each sample.

**Fig 1:**
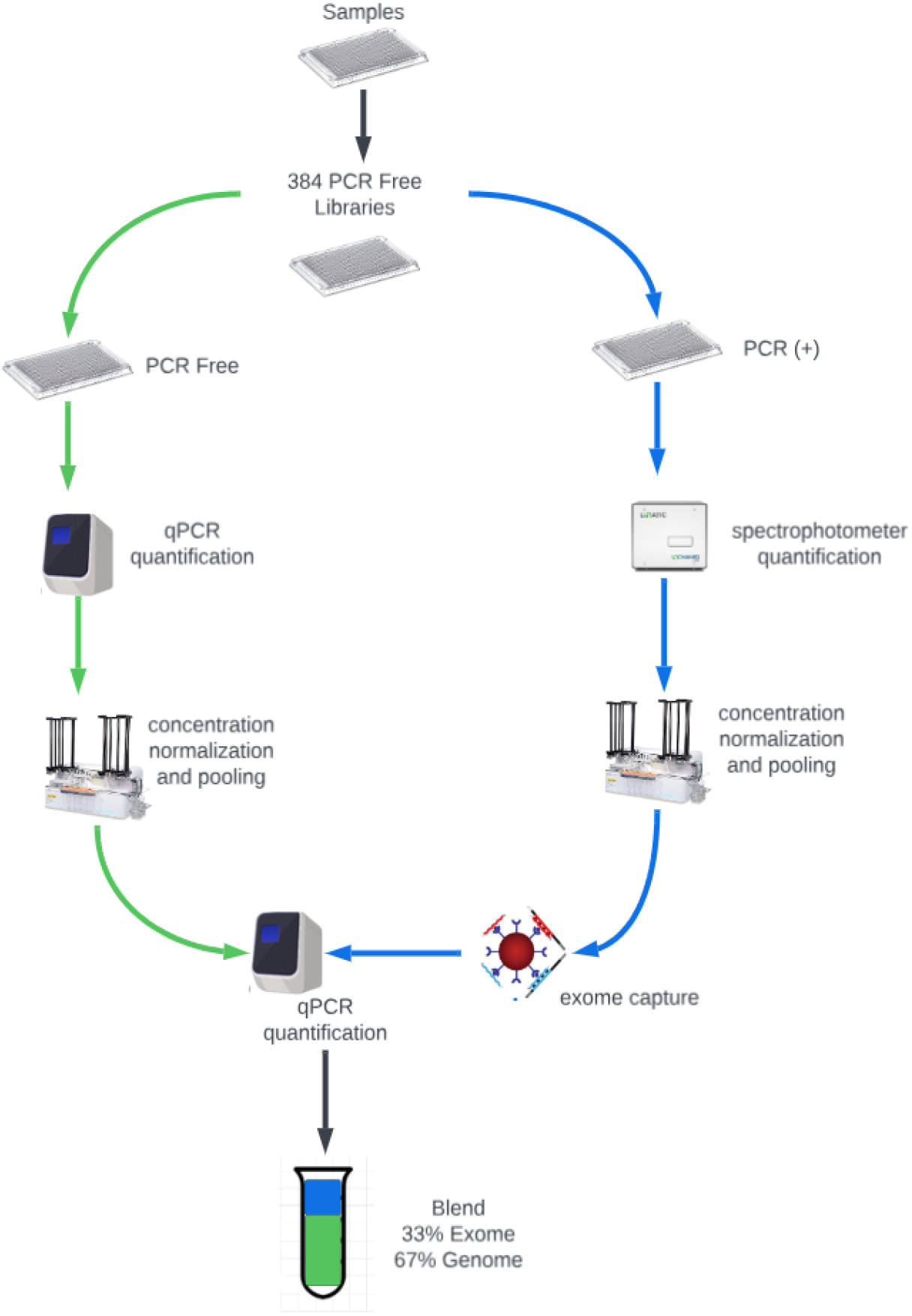
Blended Genome Exome (BGE) lab process. In the BGE lab process, PCR-free libraries are constructed from gDNA, qPCR-quantified, normalized and pooled. An aliquot of the individual PCR-free libraries is PCR amplified, and PCR+ libraries are quantified by spectrophotometer, normalized and pooled. The PCR+ library pool undergoes exome capture, is enriched and is qPCR-quantified. The PCR-free pool and the PCR+ exome captured pool are blended (33% Exome, 67% Genome) and sequenced on NovaSeqS4 (equivalent of 64 samples/lane).

Before piloting this new BGE process, we experimented to determine the best nanomolar blending ratio of whole exome to whole genome library and the amount of sequencing required per sample. Through iterative tests on samples derived from Hispanic patients (The Study of Novel Autism Genes)^7^, we titrated both the WES:WGS blending ratio and sequencing coverage to determine the best conditions for calling both coding WES and common WGS variants. Our objective was to achieve >99% r^2^ imputation concordance of BGE data to 30x whole genome data. We wanted to remain cost-efficient in sequencing, with a low coverage genome to impute the common variants, while providing enough exome coverage for 90% of exome bases to reach 10x depth. We tested, in this order, 67% WES:33% WGS for 96 samples per lane of NovaSeqS4 (Illumina, San Diego, CA, USA), 67% WES:33% WGS for 48 samples/ lane, 60% WES:40% WGS for 48 samples/ lane, 40% WES60% WGS for 48 samples/ lane, and 33% WES:67% WGS for 64 samples/ lane.

In order to analyze r^2^ imputation concordance, there were 31-62 samples among these blending ratio test samples for which we had deep whole genome data (mean 30x coverage) that could serve as a comparison truth set for concordance analysis. (Some samples were subsequently excluded during these blending ratio testing iterations as they were eventually depleted of raw DNA material for sequencing). For both BGE CRAMs and a selection of SNPs from the deep whole genomes that are present in the Infinium Global Screening Array (GSAv1.0), we used the Haplotype Reference Consortium (HRC)^8^ for phasing and imputation. Imputed calls from selected GSA SNPs were performed in RICOPILI^9^, which uses Eagle 2.3.5 for phasing and Minimac3 for imputation. We ran the BGE CRAMs through GLIMPSE^10^ version 1.1.1 for genotype refinement, phasing, and imputation. Before testing r^2^ concordance of the imputed SNPs to the deep whole genomes, we first restricted our deep genome calls only to GATK PASSing variants. Among each imputed callset, we filtered our calls to a higher quality subset of imputed calls (INFO score > 0.8 and imputed MAF difference < 15% to whole genome MAF). Per-sample concordance was then tested on all selected genotypes present in both callsets, with concordance being the squared correlation coefficient after confirming the match of locus, reference, and alternate alleles in both datasets.

The first test condition (67% WES:33% WGS and 96 samples per lane of NovaSeqS4) resulted in 29x WES and 1.5x WGS coverage per sample and >99% r^2^ imputation concordance of coding variants, but < 99% for whole genome variants. We then continued to iteratively test the other blending and sequencing coverage conditions. After plotting the varying levels of r^2^ imputation concordance (Fig 2a,b), we determined that a blend of 33% WES and 67% WGS for 64 samples/ lane provided adequate coverage of each (2-3x WGS and 30-40x WES per sample) for calling common WGS and coding WES variants (MAF > 5%), with >99% r^2^ imputation concordance to 30x WGS data for both. BGE data also provided higher r^2^ imputation concordance to 30x WGS data than imputation from selected SNPs in the Infinium Global Screening Array (Illumina) pulled from 30X WGS.

**Fig 2.**
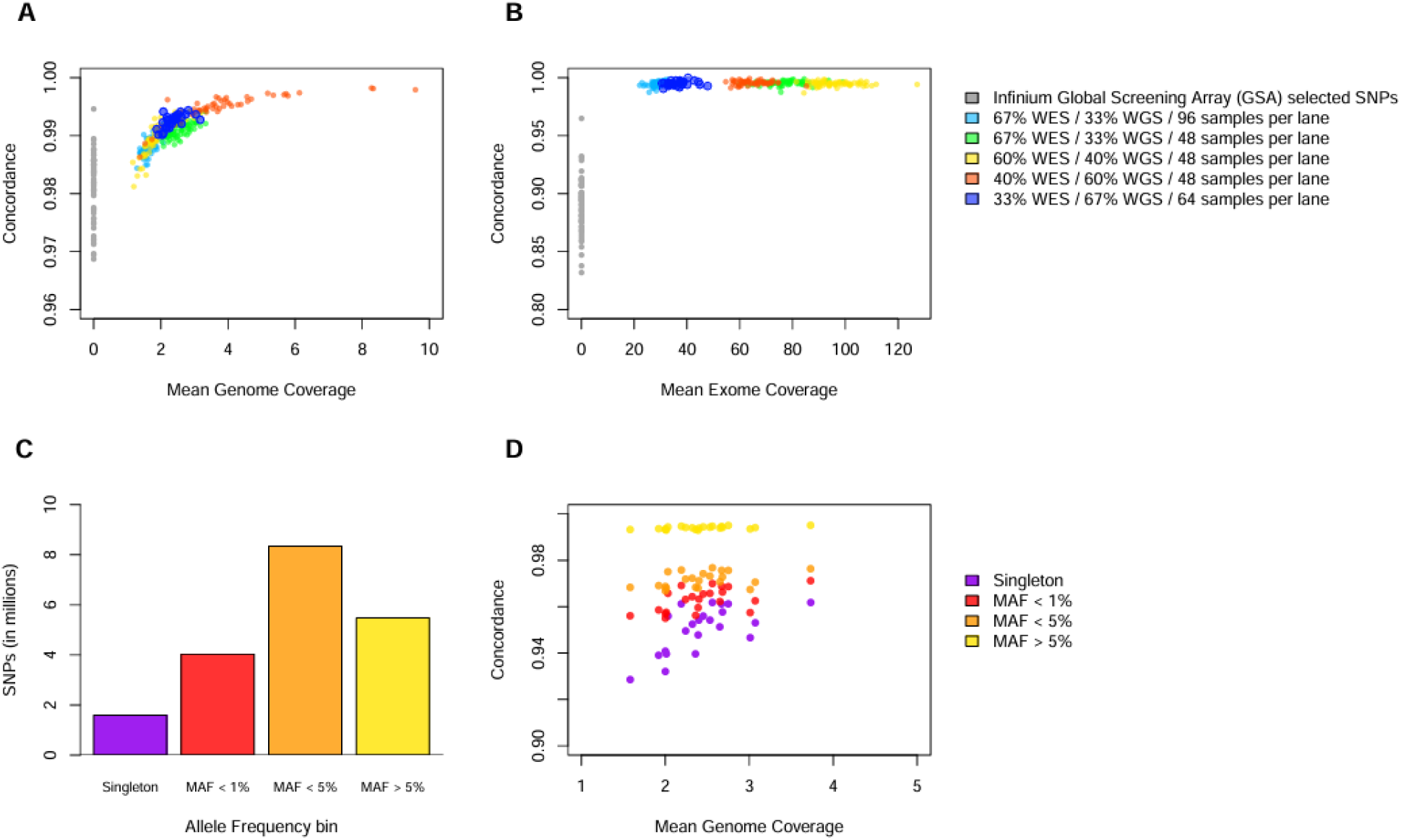
Imputation genotype concordance with deep whole genome sequencing across experiments. **a**, Per-sample genotype concordance of filtered Haplotype Reference Consortium (HRC) imputed variants to deep whole genome variants (y-axis) as a function of mean coverage (x-axis) in the low-pass genome. Gray points are not using low pass genome data, but HRC imputation results from Infinium Global Screening Array (GSA) selected SNPs from 30x whole genome sequencing. The larger dots represent the selected blending proportion and lane throughput (33% Whole Exome Sequence / 67% Whole Genome Sequence / 64 samples per lane of NovaSeq S4) for the BGE product. **b**, Per-sample genotype concordance of filtered imputed variants restricted to annotated protein-coding variants with MAF > 5% in the deep whole genome sample set (y-axis) as a function of target mean coverage (x-axis) in the exome. **c**,**d**, SNP counts (c) and concordance (d) from imputation of the full BGE pilot cohort at various allele frequency bins. Imputation was performed using the 1000 Genomes Project reference panel, and allele frequencies were estimated from 371 samples of African ancestry in the pilot cohort. Singletons are variants with only a single alternate allele genotype present in the 371 samples. Per-sample genotype concordance was done on 22 African ancestry samples with deep whole genome data.

Using the optimized blending and sequencing coverage condition (33% WES:67% WGS for 64 samples/lane), we next piloted the full BGE process, at scale, on a larger 836 sample set from blood (452) and saliva-derived (384) gDNA. These samples were derived from 64 South African, 320 Ethiopian (both from the Neuropsychiatric Genetics of African Populations - Psychosis Study and sequenced under the NIMH Populations Underrepresented in Mental illness Association Study (PUMAS)), 384 Chinese (Neuropsychiatric genetics of a Chinese population from Shanghai Province) patients, as well as 68 from the Broad Institute Study of Inflammatory Bowel Disease Genetics. Input gDNA was normalized to 50 ng/uL and plated into a 384-well PCR plate. We purified the normalized gDNA using a 2.75X solid phase reversible immobilization (SPRI) clean-up with Ampure beads (Beckman Coulter, Indianapolis, IN, USA). Purified gDNA was then quantified by spectrophotometry and re-normalized to 25 ng/uL. A target of 134 ng DNA underwent a customized fragmentation/end-repair/ A-tailing reaction for Illumina-compatible PCR-free library construction with NEBNext Ultra II FS DNA Library Preparation Kits (New England Biolabs, Ipswich MA, USA) using the following conditions: 37°C 42.57 min, 65°C 30 min. Next, we ligated unique, dual-indexed adaptors (NEBNext Unique Dual Index UMI Adaptors, New England Biolabs) to fragments (20°C 20 min) and libraries underwent two consecutive SPRI automated size-selections. Libraries were then qPCR-quantified (Kapa Library Quantification Kit, Roche, Basel, Switzerland), normalized before pooling into a single tube, and concentrated. Pooling, size-selection, and library construction were carried out in batches of 192 samples.

We took an aliquot from the pre-normalized and pre-pooled PCR-free libraries and used this as input for PCR amplification (98°C 30 sec, 12 cycles of [98°C 10 sec, 65°C 75 sec], 65°C 5 min) using the NEBNext Ultra II FS Library Preparation Kit and primers from the indexed adaptor kit (New England Biolabs). The PCR-amplified libraries were quantified by spectrophotometer, normalized to 85 ng/uL, and purified (1X SPRI). Libraries were then pooled and underwent exome capture (Human Comprehensive Exome probes from Twist Biosciences, South San Francisco, CA) using the recommended hybridization-capture protocol for xGen Hybridization Capture Core Reagents (Integrated DNA Technologies, Coralville, IA).

To calculate nanomolar concentrations for blending 33% exome with 67% genome, PCR-free and exome-captured pools were both qPCR-quantified on the same qPCR run. Blended genome-exome pools were again qPCR quantified to calculate optimal loading on sequencers. BGE pools of 192 samples were sequenced across 3 lanes of Nova S4 Lanes, using 2×150 bp reads. A single CRAM file was produced per sample.

The 836 pilot samples yielded a mean of 2.6x estimated WGS coverage and 34.1x mean WES coverage per sample. An average of 96.35% of exome bases were covered at 10x. Mean percentage of all reads aligned to the reference was 97.14 (99.5% for blood and 94.78% for saliva, with a low of 72.9% for saliva due to bacterial DNA). Twenty two African saliva samples from this pilot study had previously been whole-genome sequenced (30x), allowing us to perform the same concordance analysis as described in the earlier experiments (using the 1000 Genomes Project reference panel)^11^. We used the same filtering parameters as above, with an additional filter for genotypes with posterior probabilities < 0.9, which had a negligible impact on genotyping call rates but did increase our concordance by a few percentage points. For the larger 836 pilot sample set, we stratified our concordance analysis by allele frequency bins to see how lower frequency SNPs were performing. Allele frequencies were estimated from 371 samples of African ancestry in the pilot cohort (64 ascertained from the University of Cape Town, South Africa, and 371 ascertained from Addis Ababa University, Ethiopia). Singletons are variants with only a single alternate allele genotype present in the 371 samples, and MAF < 1% and MAF < 5% both include singletons in their frequency bin (Fig. 2c). In agreement with our blending optimization experiments, we observed >99% r^2^ imputation concordance of BGE data with 30x whole genome data, maintained high concordance over the exome, and maintained concordance levels well above 90% among singleton SNPs for these 22 samples (Fig. 2d), demonstrating the repeatability and utility of the method.

Since the original development of BGE, we have increased to processing 384 samples at a time and scaled this process to enable 300,000 samples/year. We have since applied this new method for well over 150,000 samples to provide cheaper sequencing data for imputation in large population studies. This 2-3x low-coverage genome combined with the 30-40x deep exome provides a suitable, cost-effective and unbiased improvement to SNP arrays for imputation of common variants while also cataloging rare coding variants, without any need to harmonize array data with sequencing data. This BGE method will continue to prove useful for studies of large cohorts for which deep whole genomes are not economically feasible and for studies of non-European populations for which arrays may not capture critical genetic variation. BGE, or an adaptation of BGE (different ratios of exome/genome), will also be valuable as a more affordable tool for generating polygenic and monogenic risk assessments of patients in the clinic. Rather than waiting for the cost of WGS to decrease, this method requires ten-fold less sequencing, and can immediately provide genomic researchers with many of the benefits of deep whole genomes at a fraction of the cost.

## Declarations

### Ethics approval and consent to participate

All cohorts involving human subjects that were included in this research have IRB approval following a full review of IRB applications and study consent forms by appropriate ethics committees. Details on each of the participating cohorts and IRB approvals are listed below:

1. Hispanic ASD samples, IRB title: The Study of Novel Autism Genes, Protocol **#2012p001018**, IRB of record: Mass General Brigham (previously known as PARTNERS), PI on IRB: Mark Daly
2. Chinese samples, IRB title: Neuropsychiatric genetics of a Chinese population from Shanghai Province, Protocol # **IRB17-1379**, IRB of record: Harvard T.H. Chan School of Public Health (HSPH), PI on IRB: Benjamin Neale
3. NeuroGAP-Psychosis:Ethiopia and South African samples, IRB title: Neuropsychiatric Genetics of African Populations - Psychosis (NeuroGAP-Psychosis), Protocol # **IRB17-0822**, IRB of record: Harvard T.H. Chan School of Public Health (HSPH), PI on IRB: Karestan Koenen
4. Inflammatory Bowel Disease samples, IRB title: The Broad Institute Study of Inflammatory Bowel Disease Genetics, Protocol **# 2013P002634**, IRB of record: Mass General Brigham (previously known as PARTNERS), PI on IRB: Ramnik Xavier

## Consent for publication

All participants who joined the studies listed under the above IRBs consented to the de-identified use of their data in disease research studies. During the consenting process, study staff explain the research goals and the basics of genetic processing activities. Participants are also given the option to opt out of studies if they choose. Samples used in this research and method development are all de-identified and have all been fully consented for inclusion in the study and any resulting publications.

## Availability of data and materials

Most of the data generated and analyzed during this effort were used for Research and Development only. During the pilot test, samples from the PUMAS study were included and run at production scale. These data are not publicly available yet, but per the terms of the NIMH PUMAS grant, they are scheduled to be submitted to the NIMH Data Archive (NDA). Data submission will continue once all of the data under the grant are generated and the agreed-upon embargo time has passed.

## Authors’ contributions

B.N. conceived of the project and the main conceptual idea.B.N and M.D. designed and planned the experiments. S.C. and C.S. provided project and program management support. M.D., J.G., S.D., and M.T. contributed to sample preparation and executed the experiments. M.D. and J.G. developed, optimized and scaled the customized BGE workflow. D.H, K.Y., H.H., A.M and S.Q. led the data analysis and interpretation of results. J.G. and M.D. took the lead in writing the manuscript with contributions from D.H. J.B, M.P.V, S.Q., D.S., S.T., and R.X. led studies to recruit participants, provide samples, and analyzed results for specific cohorts. All authors provided critical feedback and helped shape the manuscript.

## Acknowledgments

The authors acknowledge Scott Anderson, Andrew Bernier, Brendan Blumenstiel, Jordan Callahan, Michelle Cipicchio, Laurie Doe, Stacey Gabriel, Marissa Gildea, Kalyn Hubbard, Erin LaRoche, Matthew Lee, Atanas Mihalev, Mariela Mihaleva, Greg Nakashian, Faye Reagan, John Walsh, David Zdeb, New England Biolabs, the Broad Clinical Labs (BCL) Sequencing Team, Automation Team, R&D Team and Samples Lab. We would like to thank the patients who kindly donated samples used in these studies to make these global collaborations and resources possible to advance Autism, Bipolar, IBD and Schizophrenia genetics research.

